# Neural affect decoding from minimal spectrotemporal glimpses of voice signals

**DOI:** 10.1101/2025.07.22.663934

**Authors:** Merve Abrahamsson, Huw Swanborough, Sascha Frühholz

## Abstract

Decoding affect information from voice signals is challenging in natural environments where noise masks the most diagnostic and discriminative acoustic voice information. The extent to which neural affect decoding is resilient against or vulnerable to extreme noise-masking of voice signals remains largely unknown. Here, we performed three experiments using an auditory “bubble” technique to randomly mask and unmask certain spectrotemporal voice features similar to natural noise masking. First, using psychoacoustic voice-in-noise testing, we observed that a substantial part of the full spectrotemporal space of voice signals contains diagnostic cues essential for accurate affect decoding, contributing to the partial robustness of the voice signals under noise masking. Affective voice signals (joy, aggression) were correctly identified from early broadband spectral and temporally extended high-frequency information. Neutral voices were primarily identified from low- to medium-level spectral information with high perceptual relevance near the voice offset. Second, using human neuroimaging, we mechanistically linked neural activity to minimal spectrotemporal patches that enable accurate affect classification from voice samples. A neural network composed of (sub-)cortical auditory and limbic nodes, fronto-insular regions, and vocal motor mirroring nodes supported the affective discrimination of voice signals. More specifically, activity in the orbitofrontal cortex and amygdala was mechanistically linked to correct affect decisions, as determined by a reverse brain-to-decision mapping approach. Thus, decoding affect from voice signals embedded in complex noise seems partially feasible, provided that the noise does not obscure the minimal diagnostic spectrotemporal information essential for engaging cortical and subcortical limbic affect decoding mechanisms.

## INTRODUCTION

The ability to decode and classify voice signal information in the presence of competing noise is fundamental to our daily communication ^1^. A particular challenge faced by the human auditory system is that natural noise is spectrally, energetically, and temporally non-systematic, resulting in noise occlusion of vocal signals that are largely unpredictable by listeners. The human voice however is a distinct psychoacoustic object characterized by rich spectrotemporal features that convey both verbal and nonverbal communicative information ^2–4^, while also affording a degree of perceptual resilience to disruption from competing sounds and noise ^5,6^. This is not solely a voice general resilience, as specific vocal communication signals are encoded in acoustic channels that both carry the signal and promote further noise resilience. For example, affective voice signals demonstrate increased resilience to noise masking and recruit brain networks for socio-affective processing even at challenging threshold levels of the signal-to-noise ratio (SNR), or specifically the voice-to-noise ratio ^1^.

Despite the human auditory system’s capacity to detect and decode voice signal information at unfavorable SNRs, successful decoding still requires that acoustic features encoding this information remain perceptually accessible within unmasked spectrotemporal signal patches. The auditory system decodes stimulus global features of a partially unmasked glimpse from the perceptually available spectrotemporal information within the unmasked voice signal portion ^7,8^, analogous to the visual processing of an object’s global features despite regions being blocked from vision ^9^. Just as different visual regions of a visual object are non-informative for global feature processing (i.e. not all of the face is informative for perceiving facial affect) ^10^, not all spectrotemporal regions are equally informative for voice signal discrimination ^11^. Decoding any information carried by a partially unmasked vocal signal not only tasks the auditory brain with processing the information within the available glimpse ^12^, but also the perceptual decoding performance will be dependent on the utility and discriminative value of the information contained within those glimpses to be diagnostic for a given perceptual and/or cognitive task ^10,11,13^.

Such spectrotemporal glimpses do not have *apriori* specific acoustic profile or properties, as their perceptual and discriminative utility is rather determined by the task-relevant information they contain. This is perhaps intuitive as not only does the human voice have a specific spectrotemporal profile compared to non-voice sounds, but speech itself has a specific profile against general non-speech vocalizations ^14^. For nonverbal vocalizations, these information channels have greater importance and prominence because they are the sole source of socio-affective cues. While the above research has been essential for understanding glimpsing in human voice perception ^15^, it has largely focused on preserved features in the signal, emphasizing vocal production rather than the perceptual processes involved in vocal communication.

The aim of this study was thus to determine the perceptual and receptive side of human voice communication during the challenging condition of partial noise masking. This was achieved by utilizing previously established auditory glimpse manipulations in two main strands. First, we aimed to derive probability maps for affective voice stimuli that indicate the spectrotemporal patches, which are most relevant for the correct processing of affective information within dynamic glimpsed noise. Building on these findings, we then set out to investigate the neural mechanisms underlying the extraction of information from available spectrotemporal patches and its decoding for global affect classification and voice discrimination. Our study adapted the auditory “bubble” technique originally described for face information processing ^10^ and subsequently adapted to the auditory domain ^11^. These bubbles consist of a complete energetic mask superimposed on voice signals, with a specific ratio of the mask suppressed to zero in order to leave radial spectrotemporal profiles of the original signal detectable within the broader mask (Fig. 1).

**Figure 1.**
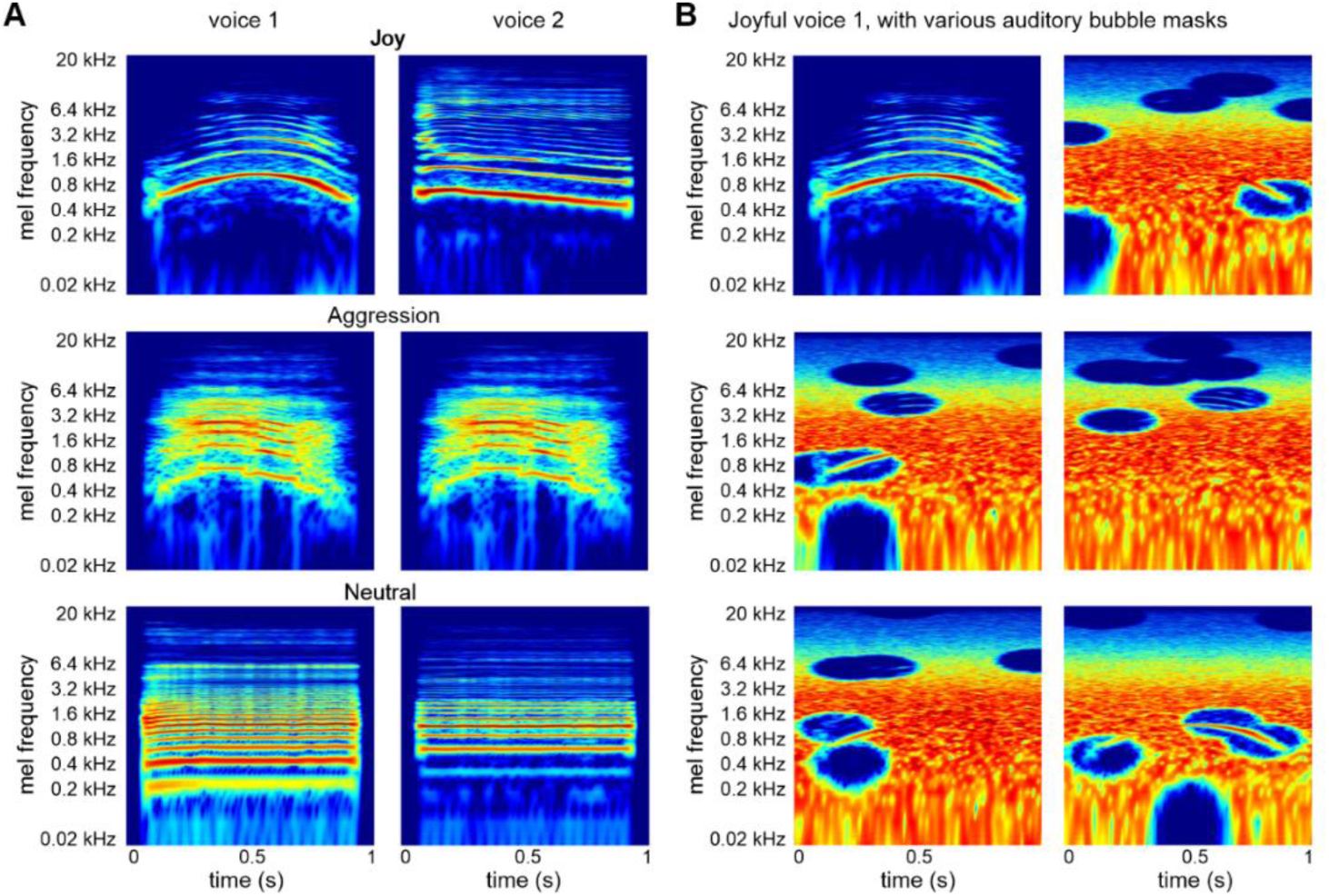
Affective voice stimuli and auditory bubble approach. **(A)** The six nonverbal affective voices were used in experiment 1 and experiment 2. Affective voices were expressed by two male speakers vocalizing an /a/ vowel in different affective intonations (joy, aggression, neutral). **(B)** One of the joyful voices is shown unmasked (top left) with various versions of the auditory bubble mask superimposed on the joyful voice sample.

Nonverbal affective voice signals were chosen for investigation as they have been previously shown to have highly distinct acoustic profiles and potentially significant spectrotemporal patches that carry relevant affect information ^16–18^. As noted in earlier work, beyond simply conferring benefits to perception in noise, affective information encoded in the voice signal elicits differential recruitment within the neural auditory system ^6^. The typical acoustic profiles of these stimuli have been described in detail for neutral voices (low fundamental frequency, clear formants and harmonics, low temporal variation), aggressive voices (higher f0, lower HNR, high energy in high-frequency ranges), and joyful voices (higher pitch, spectral variability) ^18–20^. While these features considerably contribute to affect classification and discrimination, it remains unclear to what extent they support perception under partial noise masking, and which affective voice classes may be more severely impacted by noise-related degradation.

We previously observed that affective voices are detected at lower global SNRs than neutral voices ^6^, indicating that the acoustic channels encoding affect information may confer a degree of resilience to global noise masking. This noise resilience may partly be supported by specific spectrotemporal profiles and patterns that are strongly associated with affect encoding ^21,22^ and speech intelligibility ^23^. Spectrotemporal acoustic profiles appear to be produced by the human voice in a way that is perceptually highly distinct from general non-speech background noise ^24^. Although it should be noted that high modulation power spectrum energy is often associated with increased emotional intensity, and increased vocal intensity has been shown to negatively impact affect classification tasks ^25,26^. The present study further describes the acoustic features in terms of spectrotemporal patches that meaningfully encode vocal affect, by assessing classification performance when specific spectrotemporal regions are available for the listener, adopting the same principle outlined by Mandel and colleagues^15^.

The neural mechanisms underpinning the extraction and representation of glimpsed voice information in partial and non-systematic noise are relatively unknown, though regions showing selective sensitivity for human voices are considered principal candidates. Auditory cortical (AC) regions are consistently implicated in similar tasks, demonstrating responsiveness to human voice-specific acoustic patterns ^27^, involvement in voice signal detection and discrimination tasks ^28–30^, and responsiveness to discriminative acoustic features of socio-affective voice information ^16,31–33^. In particular, superior temporal cortex (ST) regions are preferentially recruited by auditory stimuli with more harmonic structure and higher harmonic-to-noise ratio (more human voice-like), including a left-lateralized bias for human stimuli ^34^. Given that naturally occurring non-systematic noise rarely displays the same level of harmonicity and spectral variance as the human voice ^24^, this would likely be valuable in a glimpsing paradigm. Voice-like temporal features are also frequently represented in cortical regions, with the AC exhibiting noise-robust representation of temporal stimulus statistics ^35^.

Distributed regions, which are not exclusively recruited due to auditory processing, may also serve as plausible candidates for involvement. The inferior frontal cortex (IFC) is shown to be functionally involved in supporting the AC with voice signal classifications ^36–38^, and we previously showed the central role of an AC-amygdala network during voice-in-noise detection, with different network characteristics due to the socio-affective content of the voice signals ^6^. In general, the amygdala is well-established for supporting novelty and social relevance detection ^39–41^, demonstrating increased sensitivity to human over non-human voice stimuli and to spectral features encoding socio-affective information ^31^. Furthermore, the representation of voice-related temporal information has been observed in the basal ganglia system ^42^. Temporal encoding has been demonstrated along the entire auditory processing pathwayfrom the auditory nerve onwards ^43^. Cortico-limbic structures, including the hippocampus (HC), parahippocampal cortex (PHC), and insula (Ins), also exhibit entrainment to and integration of voice-specific temporal features ^44^.

While the aforementioned studies indicate involvement of multiple areas in the extraction and processing of affective voice signals and of voice-like feature sensitivity, the exact responsive networks and mechanisms underlying glimpsing and processing global stimulus features are still not known. Our study therefore addressed this specific knowledge gap using the glimpsing paradigm. Adapting previous approaches for investigating neural information sensitivity in the visual domain ^45^, we first established which spectrotemporal patches of affective voice signals are informative and discriminative during an affect classification task. Subsequently, we used a reverse correlation analysis to investigate brain regions and brain networks associated with decoding this meaningful spectrotemporal patch information. This allowed us to isolate neural responses associated with extracting brief spectrotemporal patches for stimulus-global classifications, and to examine whether these responses reflect feature processing and extraction, signal restoration and compensation, or internal perceptual template matching and integration.

## RESULTS

We experimentally mapped the relationship between spectrotemporal patches of affective voice signals that support affect identification accuracy based on minimal acoustic information (experiments 1-2). Furthermore, we aimed to examine the neural mechanisms underpinning the extraction and processing of spectrotemporal information relevant for the affect classification of voice signals (experiment 3). We utilized an auditory bubble paradigm ^15^ that produced minimal voice signal glimpses. Affective stimuli were fully masked by noise with small spectrotemporal bubbles selectively unmasking certain acoustic information (Fig. 1a-b). This allowed us to construct probability classification images of stimuli spectra from which meaningful acoustic information can be extracted in spectrotemporal patches relevant for affect classification.

### Approximating bubble ratio per time for above chance affect classification

Original stimuli were 800ms nonverbal affective voices of joyful, aggressive, and neutral tones, and all voice samples were padded with 100ms of silence at the beginning and the end of the sample. Two male speakers produced one instance of each affective voice, resulting in six voice stimuli. The stimuli were taken from a bigger database of affective voice samples and were selected for clear auditory quality and high affect recognition rates ^46^.

In experiment 1, an adaptive staircase procedure ^15^ (n=20 participants, Fig. 2a-b) was used to approximate the ability of a ∼50% performance of correct affect identification across the adaptive trials in the staircase procedure. Specifically, we varied the temporal ratio of bubbles per second in a weighted up-down staircase procedure. The spectral size of the bubbles was set to seven equivalent rectangular bandwidth (ERB) steps tall. We adapted these temporal and spectral bubble settings from a previous report ^15^. The six affective voices were randomly masked using bubbled noise with a spectrotemporal proportion of randomly positioned bubbles where noise energy was set to zero. These zero-noise bubbles exhibited a minimal proportion of the original voice signal that was not masked by noise. The information within these bubbles was thus perceptually available for auditory glimpsing of the affective information.

**Figure 2.**
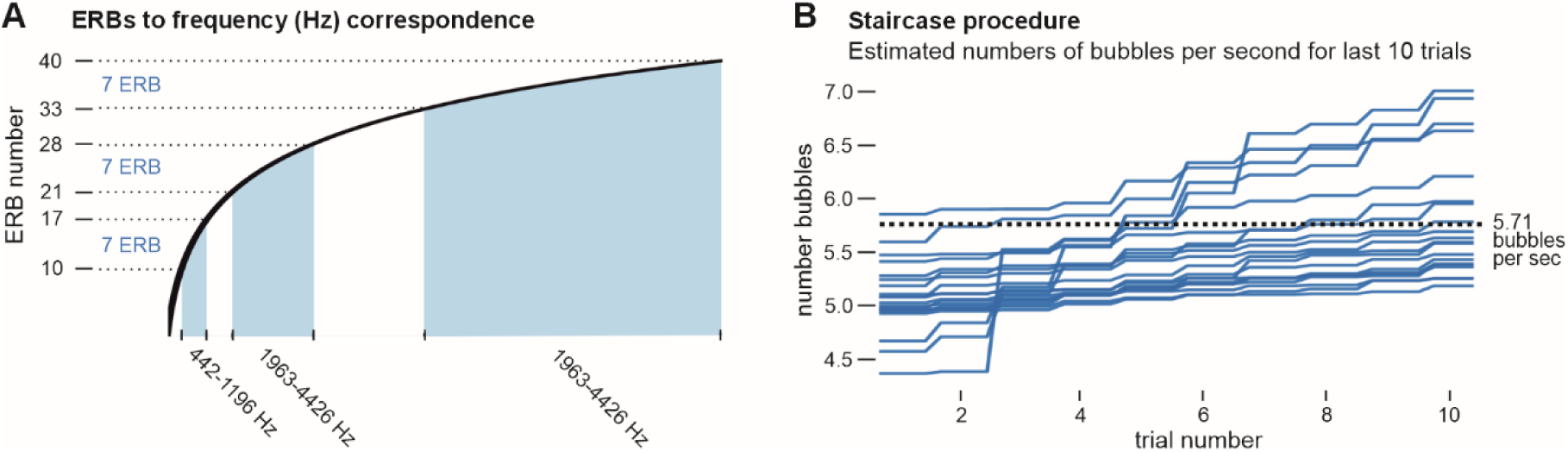
Psychoacoustic adaptive staircase experiment. **(A)** Spectrotemporal noise regions were suppressed by –64dB (relative to the noise itself) in bubbles that were seven ERBs tall (spectral dimension, y-axis) and 350ms wide (temporal dimension, not shown here) at their largest. **(B)** The bubble ratio (bubbles per second) was derived using an adaptive staircase procedure that adjusted the bubble ratio to approximately estimate ∼50% correct affect identification rate. The final bubble ratio was taken from the mean of all participants’ final ten trials.

This adaptive staircase procedure resulted in an estimated bubble size of 5.71 bubbles/sec, which approximated a ∼50% correct affect classification performance across the affective voice samples and was well above the chance level of 33%. The size of 5.71 bubbles/sec was obtained by averaging the last ten trials across participants from the staircase procedure (Fig. 2b).

### Acoustic patterns in glimpsed spectrotemporal patches for affect classification

To disentangle informative and non-informative spectrotemporal patches of voice signals that enable or disable affect classification, respectively, we then conducted a psychoacoustic experiment (experiment 2) utilizing the auditory bubble method (Fig. 1). We used the temporal size of 5.71 bubbles/sec that originated from the adaptive staircase procedure from experiment 1. While keeping the bubble ratio constant at 5.71 bubbles/sec, all stimuli were presented with equal proportions of unmasked spectrotemporal information (i.e., to avoid introducing undue influence of overall energy differences between noise masks). Each of the six affective voices was combined with 200 random auditory bubble masks. This resulted in a total of 1200 auditory samples presented randomly across six experimental blocks. As all trials in our experimental setup always contained the same amount of baseline noise derived from the full spectrotemporal profile, baseline neural activity was assumed to be non-meaningful.

Human participants (n=32) then performed an affect classification task (joyful, aggressive, or neutral) on the 1200 voice signals hidden in the partial noise. Affect classification performance across all 1200 stimuli was above the chance level with 47.27% (SEM 3.48) correct trials. Although we intended to achieve a ∼50% performance equally across all three affect categories, there were significant differences (LME analysis; F_2,38233_=78.043, p<0.001) in performance between the affect categories of joy (mean 41.55%, SEM 3.61), aggression (mean 48.27%, SEM 2.73), and neutral (mean 51.99%, SEM 4.10) (all posthoc pairwise comparisons p<0.001) (Fig. 3a). There were also significant differences between all six affective voice samples (F_2,38199_=68.368, p<0.001; all posthoc p’s<0.05), with only the neutral voice samples (p=0.879) and the Aggression 2 and Neutral 2 voice samples (p=0.326) showing no performance differences. In addition, we calculated a confusion matrix between the true affect labels and the chosen affect classes. This matrix revealed that each voice sample was classified the highest according to its true affect label, with misclassifications being lower on the alternative affect classes.

**Figure 3.**
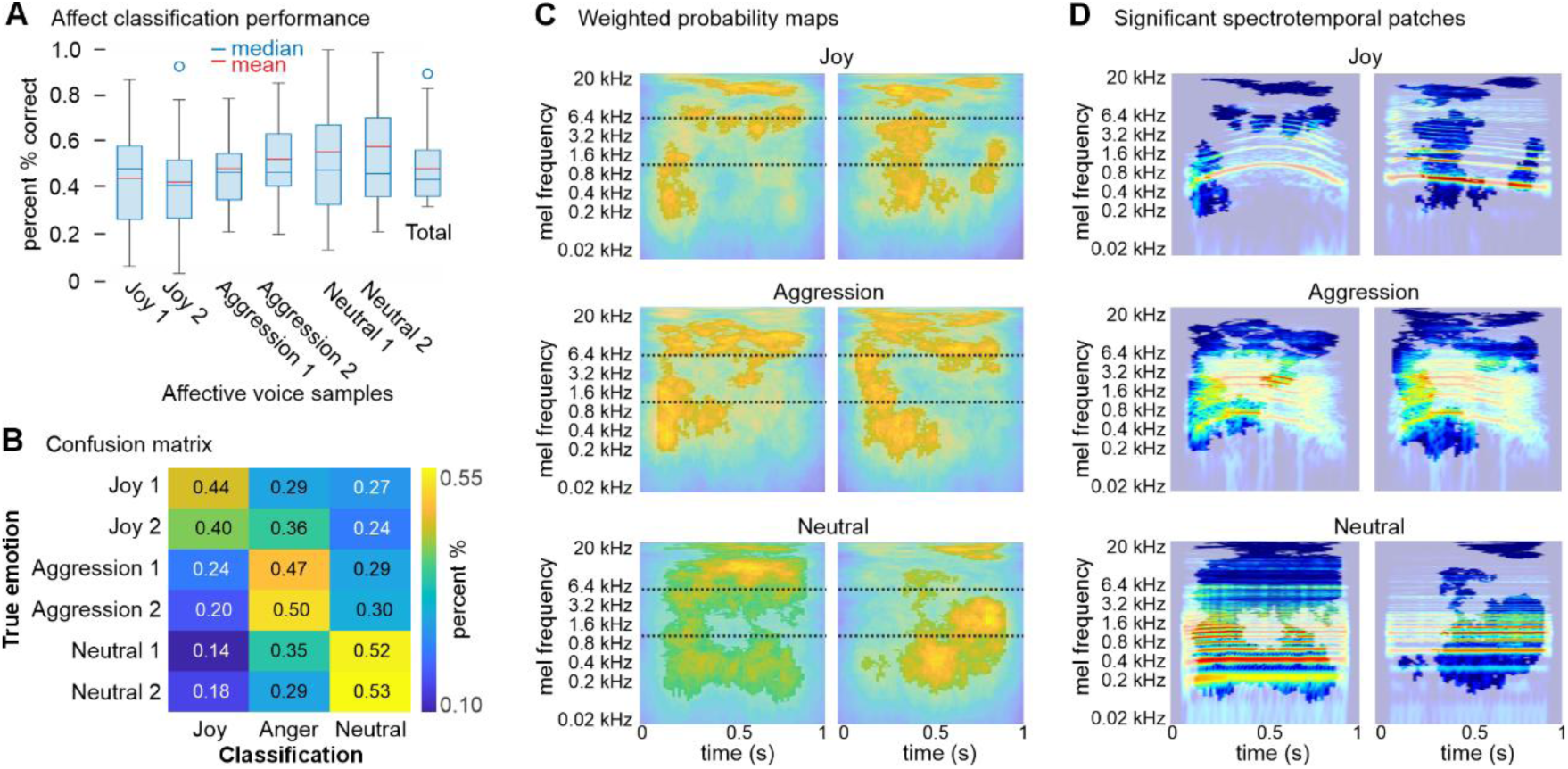
Decision patterns and significant spectrotemporal patches. **(A)** Box plots (mean, median, quartiles, and 95% confidence interval) of participant performance on affect identification of the noise-bubbled voices (200 per voice sample). **(B)** Confusion matrix for the classification performance indicating the choice probability of the affect decisions (x-axis, classification) for the affective voice samples (y-axis, true emotion). **(C)** Averaged and thresholded weighted classification maps from the binary bubble masks per participant across trials with correct affect identification. These weighted maps are entered into a group-level cluster analysis; white shading indicates non-significant spectrotemporal areas. Black dashed lines indicate frequency ranges in low (<1.6kHz), medium (1.6-6.4kHz), and high-frequency ranges (>6.4kHz). **(D)** Significant spectrotemporal patches for each voice sample and affective class overlaid on the spectrogram of each voice sample; white shading indicates non-significant spectrotemporal regions.

The affect classification performance pattern thus indicated that a substantial number of trials unmasked relevant spectrotemporal information that participants could use to make accurate affect classification. Relatedly, the remaining trials seemingly unmasked irrelevant spectrotemporal information for affect classification. The unmasked spectrotemporal information was most indicative for neutral compared to affective voice samples. This points to that spectrotemporal information for neutral voices might be more distributed across the spectrotemporal space than for affective voices. We tested this notion in the next step of the analysis.

From the classification performance, classification images were derived by aggregation of binary bubble masks used to create the noise-suppressed bubbles taken from all correctly categorized trials for each of the six voice samples. Aggregate masks were weighted by participant performance to avoid unduly influencing group effects due to outlier performance. The relevant spectrotemporal information for correct affect classification overall seemed non-random and included spectrotemporal information of high spectral power and only very minor proportions of zero-energy signal of the original voice samples (Fig. 3c-d).

More specifically, all six voice samples demonstrated significant patches of available spectrotemporal information, although the extent of the significant regions varied somewhat between stimuli and affect classes. Affective voices (joy, aggression) revealed broad significant patches across the full spectral range (low <1.6kHz, medium 1.6-6.4kHz, high-frequency range >6kHz) at the beginning of the voice samples, accompanied by a temporally extended patch in the high-frequency range (>6.4kHz). Affect voice information is often encoded and already recognized at voice onset ^47^ and in an increased spectral power of high-frequency voice information ^31,48^. The latter usually distinguishes affective from neutral voice information ^49^. Unlike affective voices, neutral voice samples revealed significant spectrotemporal patches distributed across a broad spectral range (low <1.6kHz, medium 1.6-6.4kHz, high >6.4kHz), but partially more extended in the temporal dimensions. There was especially a temporal weighting in the second half of the voice samples, predominantly in the lower frequency ranges. Classifying voice samples as neutral often requires decoding acoustic information across the full temporal range to confidently exclude the presence of any potential affect information over time. Classifying voice signals as neutral might also shift attention to low-frequency ranges, as they contain the predominant spectral information in neutral voices^49^.

### Neural patterns supporting affective voice glimpsing

In experiment 3, we investigated the neural mechanisms of glimpsing affective voice information and the brain mechanisms implicated in the extraction and processing of the spectrotemporal patch information that facilitates affect classification. A new cohort of human participants (n=24) listened to the same 1200 trials and performed an identical affect classification task as in experiment 2 during functional magnetic resonance imaging. Our analysis followed three approaches as described below, with the overarching goal of differentiating the neural activity responsive to the task and the neural activity responsive to the assumedly meaningful spectrotemporal patches.

First, because the primate auditory cortex contains specific fields for the processing of voice features and voice patterns ^28^, a functional voice localizer scan was performed. This allowed us to derive a comparative activation map of the cortical voice area (VA), representing voice-selective brain regions in the superior temporal cortex (ST). The functional definition of the VA was used to potentially localize neural activity related to voice features and voice pattern processing during the main experiment in dedicated cortical voice-sensitive areas ^27,28^. Comparing the neural response to voice sounds to that to the animal and environmental sounds, we found extended auditory cortical activity covering areas in Heschl’s gyrus (HG), planum temporal (PTe), and higher-order auditory cortex (Te3) (Fig. 4a, Tab. S1).

**Figure 4.**
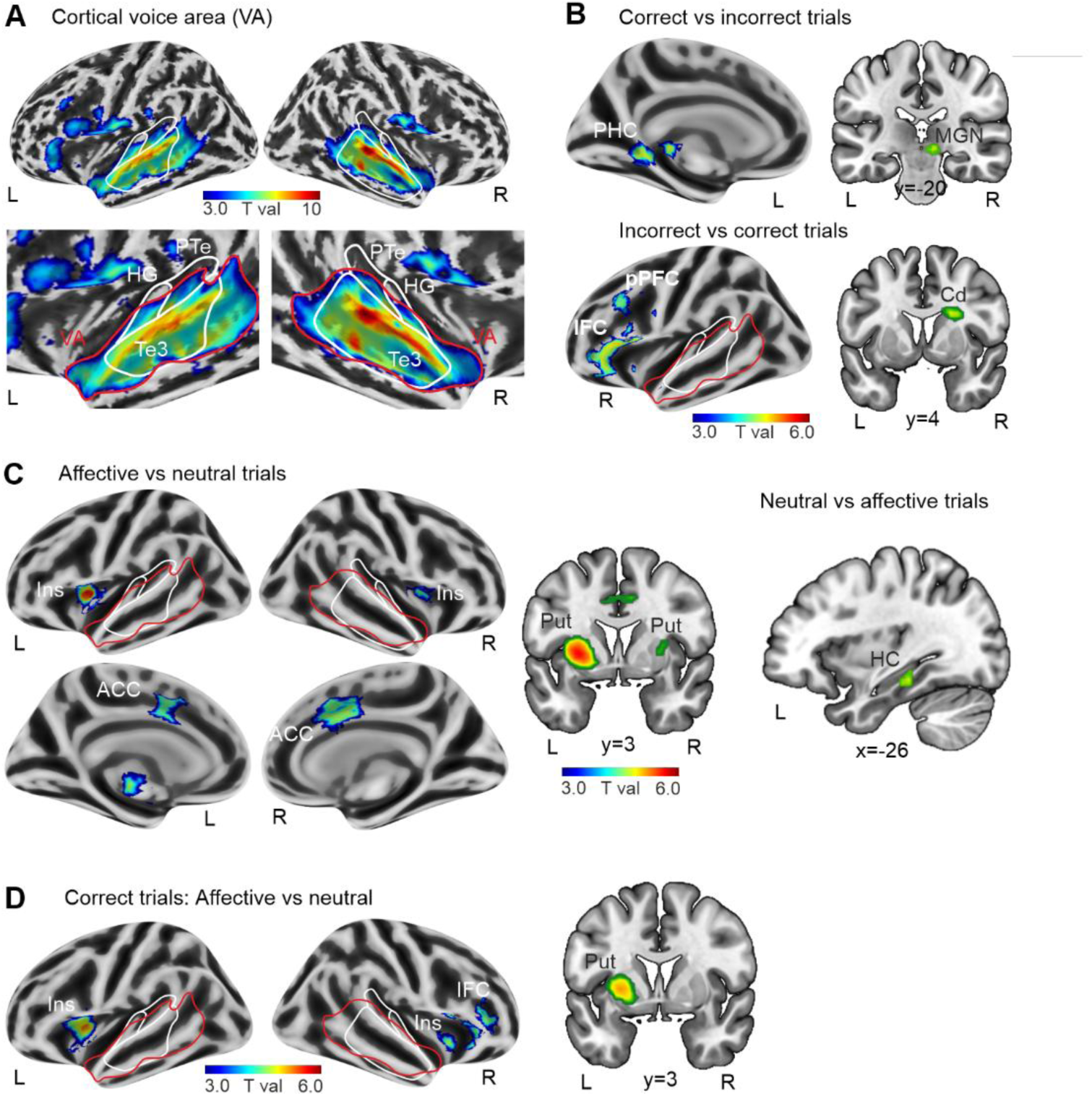
Brain activity for comparing experimental conditions. **(A)** The cortical voice area (VA), as determined by a separate functional localizer scan, shows extended bilateral STC activity across auditory cortical regions (*HG* Heschl’s gyrus, *PTe* planum temporale, *Te3* higher-order auditory cortex). **(B)** Functional brain activity for contrasting correct and incorrect trials, with the contrast [correct > incorrect] revealing activity in the fusiform gyrus (FG) and medial geniculate nucleus (MGN) of the thalamus; the reverse contrast [incorrect > correct] revealed activity in the inferior frontal cortex (IFC), posterior prefrontal cortex (PFC) and caudate nucleus (Cd). **(C)** Functional brain activity for contrasting affective with neutral trials, with the contrast [affect > neutral] revealing activity in bilateral insula (Ins), anterior cingulate cortex (ACC), and putamen (Put); the reverse contrast [neutral > affect] revealed activity in left hippocampus (HC). **(D)** Functional brain activity for contrasting affective and neutral trials only for trials with correct affect classification, which revealed activity in bilateral Ins and right IFC, and left Put. Combined voxel threshold p<0.005 and cluster size threshold k=31, resulting in p<0.05 corrected at the cluster level.

In the second analysis approach, we first compared brain activity for trials with correct classifications to that for trials with incorrect classifications to examine the neural activity that influenced the accuracy of affect classifications. Correct affect classification (Fig. 4b) was associated with activity in the medial geniculate nucleus (MGN) as part of the ascending auditory pathway to perform basic spectrotemporal acoustic analysis in the unmasked patches ^50,51^. Further activity was found in the parahippocampal cortex, potentially matching incoming sounds to stored representations ^52^ that might support the perceptual pattern completion of degraded sound ^52^. Incorrect affect classifications on noise-masked voice samples were associated with neural activity in the prefrontal cortex (PFC), inferior frontal cortex (IFC), and in the right caudate nucleus (Cd) of the basal ganglia complex. Frontal regions are typically associated with classifying communication sounds ^53^, while the caudate can set and shift the decision criteria between classification options ^54^. Since no feedback was given to participants, activity in the PFC might represent a level of subjective confidence in the accuracy of the decision ^55^.

Neutral and affective voices have different spectrotemporal profiles and thus might evoke different brain activity, even in the case of partial noise masking. Affective trials evoked higher neural activity in the insula (Ins) and anterior cingulate cortex (ACC) which typically perform an external sensory to the internal physiological mapping of perceived affective signals ^56^ as well as in putamen (Put) for sensory gating ^57^ (Fig. 4c). Neutral trials showed higher activity in the hippocampus (HC), presumably to retrieve associative and episodic information ^58^ for assessing a voice signal as being of neutral or non-affective quality in noisy conditions.

We also compared neural activity for affective trials to that for neutral trials, conditional on the correctly classified voice samples (Fig. 4d). A similar neural pattern was revealed for the contrast including all trials (Fig. 4c), but with additional activity in the right IFC. Unlike the left IFC, which was found for incorrect trials, the right IFC is essential specifically for affect classification in voice signals ^53^ and might be a neural indicator for performance accuracy in affect decoding from voice signals under challenging conditions. This might point to a right-to-left gradient in IFC for coding the explicit and implicit accuracy of affect classification performance.

### Reverse mapping of brain patterns to decision-relevant spectrotemporal patterns

In the third analysis approach, we aimed to establish a closer association between the spectrotemporal patches that contained relevant voice signal information and brain activity. Hence, we performed a reverse correlation analysis to derive the neural responses associated with the perceptual availability of the meaningful spectrotemporal information for each stimulus^45^ (Fig. 5a-b). For every subject and voxel, voxel responsiveness to spectrotemporal bubbles was assessed, similarly to the classification images in the psychoacoustic experiment (experiment 2). Each voxel was split at the median point for high-responding and low-responding trials according to their beta weight for each of the 1200 trials. Binary bubble masks for high- and low-responding trials were, respectively, aggregated into positive (mean of top 50%) and negative response maps per voxel (mean of bottom 50%). Next, the negative response maps were subtracted from the positive response maps, resulting in a difference map representative of the spectrotemporal patches to which the voxel was maximally responsive (Fig. 5a). These difference maps were then correlated with the six individual stimuli response maps derived from the psychoacoustic experiment 2 (Fig. 3c). This yielded six whole brain images of neural activity associated with spectrotemporal regions carrying relevant information for affect classifications per subject (Fig. 5b).

**Figure 5.**
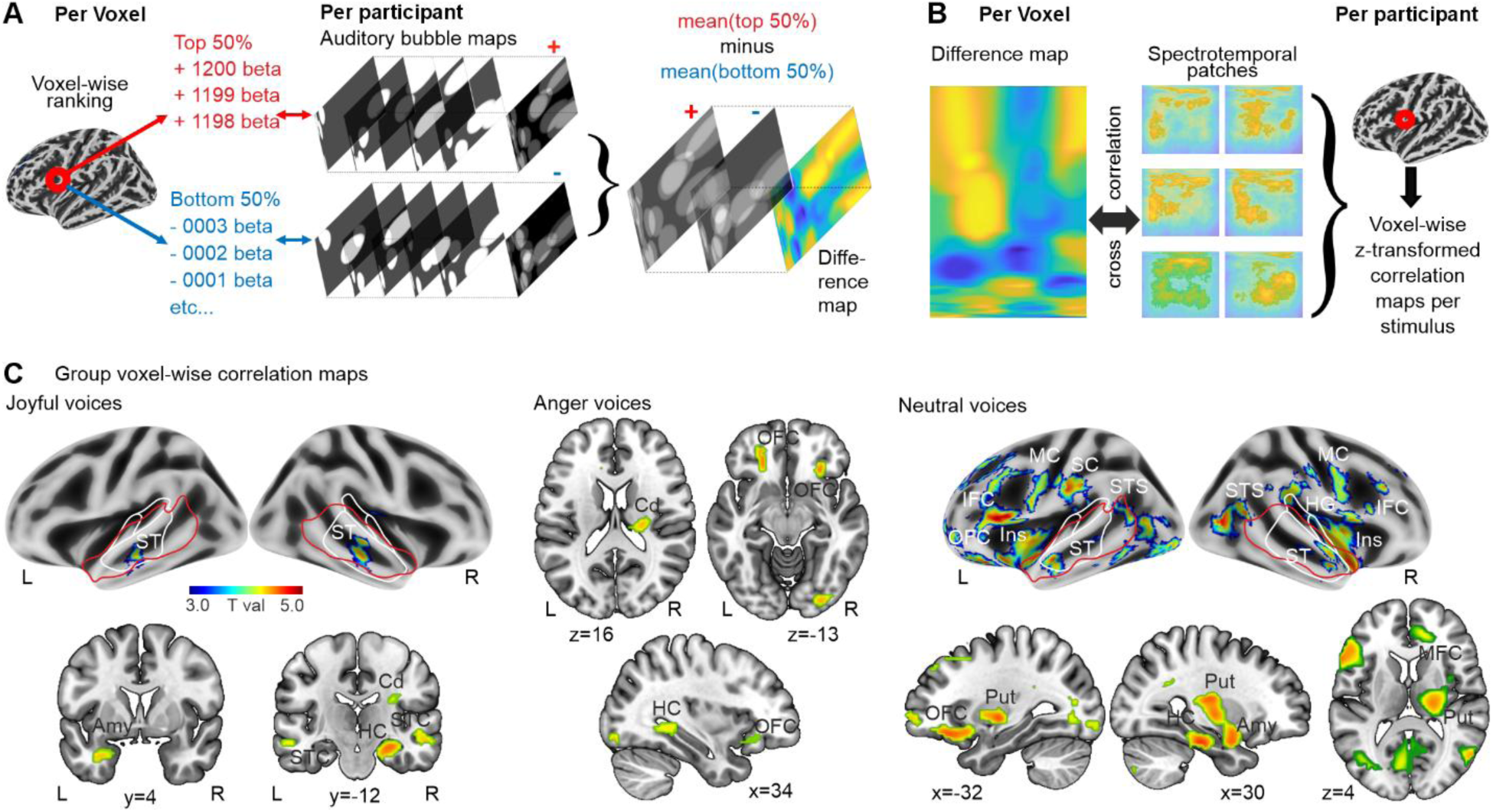
Reverse correlation analysis. **(A)** Finite impulse response functions (FIR) were used to model each presentation of the 1200 individual trials. For each voxel, the beta weights are z-scored and median-split as high (red) or low responses (blue) for an individual voxel. Mirroring the analysis from the psychoacoustic experiment, statistical maps are created by summing the bubble masks for each trial according to their being high (mean of top 50%) or low response (mean of bottom 50%) per individual stimulus. The masks made from low-responding trials are then subtracted from the high-response masks to create a spectrotemporal heatmap for each individual voxel. **(B)** Each difference per original stimuli is correlated with the significant image classification map for its relative stimuli, creating brain volumes of correlation maps. **(C)** Group-level correlation maps separately for joyful (left panel), aggressive (mid panel), and neutral voice samples (right panel). Voxel threshold p<0.005, cluster size threshold k=31, combined p<0.05 corrected at the cluster level.

Group-level voxel-wise correlation maps revealed significant brain-to-behavior associations with some commonalities but mostly differential results for the three vocal affect classes (Fig. 5c, Tab. S2). Vocal joy revealed significant brain-to-behavior associations in the right Cd and right HC as well as in the left amygdala in bilateral ST regions that were located within the voice-sensitive AC. The latter two regions are core nodes in the neural affective voice processing network to integrate voice patterns and affective information for stimulus evaluations ^31,56^. Similarly, aggressive voices revealed significant brain-to-behavior associations in the Cd and HC accompanied by unique activity in the bilateral orbitofrontal cortex (OFC), potentially to gauge the social significance of the aggressive voices or voice features that indicate social aggression ^51^. Compared to the affective voices, neutral voices revealed the most extended behavior-to-brain associations across cortical and subcortical regions. Significant effects were found in the limbic system (Amy, HC), in ST inside and outside the voice area, the putamen (Put), medial frontal cortex (MFC), the fronto-insular system (IFC, OFC, Ins), and the inferior sensory-motor system (MC motor cortex, SC somatosensory cortex). Although this network of regions is typically involved in decoding affect information from sound and sound features using acoustic ^28^, social ^51^, associative ^58^, and motor mirroring analyses ^59,60^, it might also help to inversely determine the absence of affective sound features in the case of neutral voices.

### Mechanistic brain patterns that support correct affect classifications

The reverse correlation analysis revealed brain areas associated with the assumed availability of meaningful spectrotemporal patches within stimuli during the auditory glimpsing of voice samples of three affect classes. To obtain a more differential picture of the involvement of certain neural nodes that support correct affect classifications, we further analyzed the brain-to-behavior association patterns. We first performed a whole-brain analysis to identify brain regions that showed common effects of brain-to-behavior patterns across all three affective voice classes (Fig. 6a, Tab. S3). A conjunction analysis identified the left amygdala and the right putamen as neural nodes that statistically showed common effects across all voice samples. Second, in a further analysis using group-level F-contrasts, we determined neural nodes that show differences between the affective voice classes (Fig. 6b, Tab. S3). This analysis identified significant effects in the left IFC, OFC, Ins, and Put. These common and differential neural nodes might be associated with more granular processing for decoding spectrotemporal patch information that potentially determines both the general and specific accuracy of affect classifications.

**Figure 6.**
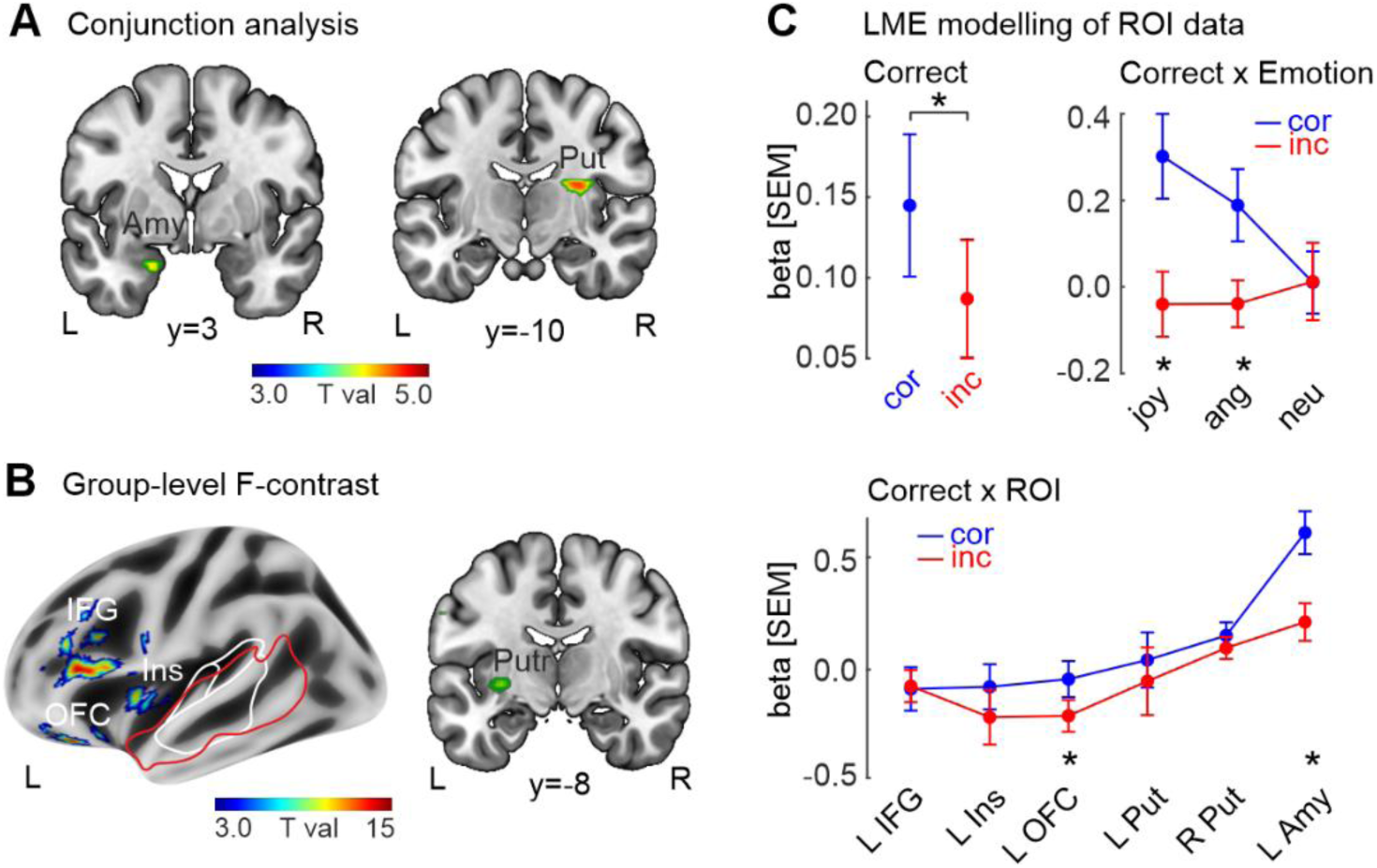
Correlation maps and ROI data patterns associated with correct affect classifications. **(A)** Group-level conjunction analysis across all correlation maps (p<0.005, k>31; p=0.05 corrected at cluster level) with peak activity in the left amygdala (Amy) and right putamen (Put). **(B)** Group-level conjunction F-contrasts across all correlation maps (p<0.005, k>31; p=0.05 corrected at cluster level) with peak activity in left orbitofrontal cortex (OFC), inferior frontal gyrus (IFG), insula (ins), and putamen (Put). **(C)** Estimated effects of experimental factors on single-trial beta-weights from regions of interest (ROI) defined by the conjunction analysis and the F-contrast. The LME analysis indicated a main effect of the factor correctness, with higher ROI activity for correct compared with incorrect trials (left upper panel). The factor correctness also interacted with the factor affective voice class (right upper panel), with higher brain activity for correct joy and anger trials. There was also an interaction between the factor correctness and the factor ROI, with higher brain activity for correct trials in the left OFC and left amygdala (lower panel). Asterisks * indicate significant differences between correct and incorrect trials at p<0.05.

To determine the relevance of activity in these neural nodes for correct affect classifications, we extracted the local signal in these regions of interest (ROI) in the 1200 single-trial beta-weight images. An LME analysis was used to determine the significance of the factor correctness (correct trials, incorrect trials) for the signal patterns in the ROIs. First, we found an overall influence of the factor correctness on the signal of the ROI (F_1,182890_=8.204, p<0.001), with correct trials showing a higher signal than incorrect trials. Second, there was an interaction effect between the factor correctness and the factor affective voice class (F_2,183498_=4.013, p=0.018). Specifically, vocal joy (p<0.001, FDR corrected) and aggression (p=0.003 FDR) showed a significant difference in the ROI signals between correct and incorrect trials but not neutral voice samples (p=0.847 FDR). Third, the factor correctness also revealed an interaction effect with the factor ROI (F_5,184102_=3.528, p=0.003). Significantly higher signals were observed only in the left OFC (p=0.004 FDR) and left Amy (p<0.001 FDR) during correct trials as opposed to the incorrect ones. Taken together, it seems that a specific cortical and subcortical limbic network supports the correct classification especially of affective voices that are glimpsed from limited acoustic information in meaningful spectrotemporal patches of the voice signal.

## DISCUSSION

Glimpsing affective information from considerably noise-masked voice signals seems possible, but this depends on specific features of the unmasked spectrotemporal cues and underlying neural mechanisms. We here experimentally investigated an ecological scenario in which natural noise is spectrally, energetically, and temporally non-systematic, posing challenges for the accurate identification and classification of socially relevant affective voice signals in natural settings.

Using an auditory bubble technique, we first identified spectrotemporal patches that reliably support accurate affect classifications of voice signals, even if substantial portions of the signals are obscured by noise. There were three major findings for the properties of the spectrotemporal patches across the affective voice samples. First, there were some critical differences between affective and neutral voices. Affective voices (joy, aggression) could be identified by broadband spectral range of voice information relatively early at the onset of the voice signal with additional support by a temporally broad patch of high-frequency information across the duration of the voice signal. The presence of affective information on voice signals can be very rapid ^47^, as basic affective voice signals already show a clear acoustic difference to neutral voice at voice onset ^48^. Furthermore, affective voices usually have higher energy in medium and high-frequency ranges of the voice signal ^49^, which sets them apart from neutral voices. Second, although vocal joy and vocal aggression revealed a similar overall pattern of spectrotemporal patches, there were also distinct differences that may have facilitated the discrimination of these affective tones. Namely, high-frequency patches were relatively larger for aggressive voices, and the low-frequency patch for aggressive voices extended temporally from the onset into roughly the midsection of voice samples. Aggressive voices usually have a distinct lower harmonic-to-noise ratio as well as relative increased power of higher compared to lower frequencies, which is referred to as the Hammarberg index ^49^. Both of these features could be computed from the larger high-frequency and low-frequency patches. Compared to aggressive voices, joyful voices revealed a small low/medium patch towards the voice offset, especially for one of the joyful voice samples. This patch unmasked a distinct spectral profile, including the fundamental frequency and several voice formants, and thus appeared like a final confirmation of the pitch profile that is commonly observed for joyful voices ^31,49^. Third, and finally, neutral voices revealed a temporally extended patch in the lower frequency band and a broader low/medium patch near the voice offset. The latter patch might be a confirmation patch that reflects the full presence of a neutral voice, while it may concurrently imply the absence of any affective information until the offset of the voice signals. The patch in the high-frequency range may further suggest this second interpretation. Neutral voices usually do not have energy in the high-frequency range, and the absence of such high-frequency information might signify the neutrality of a voice signal ^48,49^.

As indicated by the spectrotemporal patch analysis, voice signals can be correctly classified based on minimal acoustic information glimpsed from the spectrotemporal properties of voice samples. This analysis further revealed that a large portion of the spectrotemporal voice information seems negligible or potentially irrelevant for making such affective classification, at least during conditions of isolated glimpsing into specific spectrotemporal patches. This is an interesting observation, since much of this seemingly irrelevant information usually contributes to the acoustic description of affective voice profiles ^46,49^. This highlights the notion that not all acoustic features of affective voices are perceptually relevant for listeners, especially when making affect decisions under challenging listening conditions.

Such affect decisions during minimal voice glimpsing are supported by specific neural mechanisms in a distributed brain network. Affect decoding from voice signals typically involves a central network of limbic, auditory, and fronto-insular brain regions ^14,56^. Beyond the activity in several of these regions of the affective voice processing network, we have also observed the neural contribution of additional brain areas that facilitated the challenging task of affect decoding from minimal acoustic input. Correctly identifying the affective tone from glimpsed voice information involved the MGN as a pre-cortical node for spectrotemporal pattern analysis ^50,51^ and the PHC supporting pattern completion in case of incomplete sensory information ^52,61^. More specifically, correct trials for affective voices involved a network of bilateral insula, putamen, and right IFC. The putamen could provide sensory gating mechanisms ^57^, which the insula could use for mapping sensory information to internal affect representations ^56^. Based on this sensory analysis and affect decoding, the IFC is commonly implicated in the classification of communication-relevant sounds ^62,63^. For correctly classified affective voice signals, we observed a specific involvement of the right IFC. This provides a specific affect representation based on the voice quality profile and the specific pattern of features ^53^.

In contrast to neural activity in right IFC for correct trials, incorrect trials were associated with activity in left IFC together with left PFC activity. This points to a frontal hemispheric asymmetry regarding the distinction between the objective and subjective level of correctness of affect classifications. While the right IFC encoded the objective correctness of the affect decision, the left IFC was active during incorrect trials. Participants in our experiments did not receive feedback on the accuracy of their affect classification performance. As a consequence, they might have responded incorrectly in trials with highly ambiguous or irrelevant spectrotemporal information unmasked by bubbles. Such responses might also entail heightened levels of uncertainty during incorrect trials. While the left IFC might perform probabilistic predictions about the likely affect signal ^64^, the left PFC may encode the level of uncertainty associated with such a decision ^65^.

To further explore how neural activity mechanistically relates to spectrotemporal patch information that generally and specifically support accurate affect classification, we performed a reverse brain-to-behavior mapping approach in two steps. We first linked single-trial neural activity with the significant spectrotemporal patch information for each affective voice class. Joyful voices revealed a brain-to-behavior mapping in an auditory-limbic network, including the amygdala. This represents the core neural network for affective sound analysis ^31,66^. Aggressive voices exhibited a unique contribution of the OFC as a neural node for social evaluation of affective stimuli ^67^, which is especially relevant for vocal aggression ^68^. Neutral voices showed a largely extended cortical and subcortical network for this brain-to-behavior mapping, including neural nodes (auditory, limbic, OFC) that were also found for the affective voices. Further contributions were revealed by the MFC and the inferior somato-motor cortex, which might provide an in-depth evaluation ^56,69^ and motor mirroring for voice feature analysis^70^. Neutral voices had the largest spectrotemporal patches uncovered, reflecting a high amount of acoustic information for correct neutral classification. The acoustic information in neutral voices requires more elaborate neural processing, given that decisions about the neutrality of a voice sample are often more challenging and are frequently performed by eliminating the affective options ^51^.

While this first step of the brain-to-behavior mapping analysis linked neural activity to significant spectrotemporal patch information, in the second step, we further linked neural activity to the correctness of affect decisions. This step uncovered that especially neural signals in the left OFC and the left amygdala were significantly higher during correct than incorrect trials in the affect classification task. Thus, in contexts involving minimal voice signal glimpsing, neural mechanisms involved in the social and affect analysis of partial acoustic information support the accurate decoding and classification of affect information ^56^. Surprisingly, we did not find a significant link of subcortical and cortical auditory nodes that linked neural signals to the accuracy of affect decisions. The subcortical MGN showed statistically significant activity during the original analysis when contrasting correct with incorrect trials. The cortical ST showed a significant brain-to-behavior mapping for joyful and neutral voices when linking neural signals to the significant spectrotemporal patches. It might be that auditory neural nodes contribute to preparatory acoustical analyses that enable socio-affective classification processs within the OFC-amygdala network.

Taken together, the human neural system can partially classify affective content on minimal acoustic information under substantial noise masking, which can be of critical social and survival relevance in certain contexts. However, such affect decoding from glimpsed voice signals critically depends on the absence of certain noise properties that would otherwise occlude relevant affective voice information to the point of being unrecognizable.

## MATERIALS AND METHODS

We conducted three experiments with independent samples of participants. All participants had normal hearing with normal/corrected-to-normal vision and had no history of neurological or psychiatric disorders. All participants provided written informed consent in accordance with the ethical and data security guidelines of the University of Zurich and of the Zurich Cantonal Ethics Commission. Ethical approval for all three experiments was obtained prior to data collection.

### Experimental stimuli

The stimuli set for all three experiments was identical. We selected six original recordings of nonverbal affective expressions from a voice dataset that has been used in previous studies ^2,46^. In these recordings, two male speakers each produced vocalizations in three affective tones (joy, anger, neutral). All six vocalizations were produced on the vowel /a/ and had a fixed duration of 800 ms (Fig. 1a). All vocalizations were RMS normalized and were scaled to 70 dB SPL. Each voice sample was padded with 100 ms silence before and after the sample, resulting in 1000 ms sound samples. These samples were each embedded in 200 random mixes of bubbled noise that selectively preserved spectrotemporal regions of the target voices whilst fully masking the rest of the stimuli (Fig. 1b). As the goal of the study was to assess the spectrotemporal regions that facilitate successful voice glimpsing, we opted to not include both female and male speakers, because this would have extremely extended the study and would not have allowed to accurately account for the differences in the voice fundamental frequency between sexes.

When creating the bubbled noise mixes, we consulted previous work on the measurement of specific spectrotemporal regions for speech comprehension in noise, including the parameters for bubble size, number of mixes required for adequate stimuli masking, and noise generation ^11^. The noise acoustic profile (prior to having bubbles added) was generated by performing a short-time Fourier Transform (STFT) of the concatenated vocal stimuli (0.25 ms bins) and taking the 97^th^ quantile for each STFT band’s amplitude. This was multiplied across the columns of the STFT of the white noise (0.25 s bins), re-weighting the white noise to 97^th^ quantile of vocal stimuli frequency magnitudes. The signal was reconstructed with an inverse STFT with a constant phase shift applied across all bands.

Voice-plus-noise mixes were adjusted so that voice samples were completely undetectable when no bubbles were applied to the mask (-24 dB SNR). For each voice-plus-noise mix, bubble masks were created that suppressed the general noise by -64 dB in the randomly generated spectrotemporal bubble regions. This procedure ensured that the acoustic content of the original voice samples in any given bubble region would be undisrupted by masking noise, while all other acoustic information of vocalizations was masked by the noise.

### Staircase estimation of auditory bubble size (experiment 1)

We recruited a sample of 20 healthy participants (13 female, mean age 23.6y, *SD* 3.9) to take part in a staircase experiment. Here we sought to estimate the required bubbles per second (Fig. 2a-b) to achieve noise-stimuli mixes that would be at approximately a 50% performance (i.e., such that approximately 50% of trials would be detectable). The details of this staircase assessment are described in Mandel and colleagues ^15^.

Experiment 1 comprised of 75 total trials with varying number of bubbles per second. The experimental procedure started with nine bubbles per second, followed by a weighted up/down staircase procedure by varying the number of bubbles per second. The spectral size of the bubbles was fixed to seven equivalent rectangular bandwidth (ERB) steps tall. The experiment was conducted in an anechoic chamber, using Sennheiser® HD 280 Pro headphones for stimulus presentation.

The threshold performance in the psychoacoustic staircase assessment was estimated to be 5.71 bubbles per second. All parameters for bubble sizes and generation were left unchanged from the original paper ^11^. Bubble center locations were selected uniformly at random locations in time and ERB space, excluded from the two ERBs at the top and bottom of frequency space. At their maximum height and width, the bubbles were seven ERBs tall and 350 ms wide (Fig. 1b).

### Psychoacoustic experiment (experiment 2)

Experiment 2 aimed to establish spectrotemporal patches of the stimuli important for accurately identifying affective tone in voice samples, following the methods established by Mandel and colleagues ^11^. To achieve this aim, we recruited 32 participants in this experiment (19 female, mean age 24.3y, *SD* 4.2). Participants were recruited via public advertising at the University of Zurich. The experiment was conducted in an anechoic chamber, using Sennheiser® HD 280 Pro headphones for stimulus presentation.

The experiment was carried out in a single session, with a total 1200 trials divided across six blocks such that 200 mixes of each vocalization presented randomly across the experiment. Stimuli were randomized across all stimuli conditions (joy, anger, neutral). A fixation cross was displayed at the center of the screen at the start of each trial and remained on the screen for the entire trial. In each trial, participants listened to a single voice-plus-noise mix and they were asked to classify affective tone (joy, anger, neutral) of the voice via button presses using their right hand. Each trial was separated by an intertrial interval of 2-5 s. The order of buttons was balanced across all participants in both studies. No feedback was provided to participants.

### Statistical estimation of spectrotemporal patches of important affect information

All psychoacoustic analysis was conducted using MATLAB 2021b. Across all participants, the binary noise-bubble masks (1 bubble area, 0 noise mask area) were summed together for all trials with a correct affect classification for each of the six stimuli. The masks were weighted for each participant as the inverse of that participant’s total accuracy, preventing high performers from unduly influencing the analysis. This procedure resulted in a corresponding response map for each stimulus, which quantifies the probability of local spectrotemporal information in the voice samples that most likely contributes to accuracy in affect classification.

Statistical testing of these response maps was performed using the random field theory cluster analysis toolbox Stat4Ci ^71^. This toolbox is designed for statistical testing of classification images lacking *apriori* assumptions, particularly when using bubbled stimuli and reverse correlation paradigms. Within each image, Stat4Ci computes the likelihood of observing a cluster of z-values in clusters of *k* size or greater above a given threshold (*t)*. Cluster sizes are variable depending on z-values in the coherent cluster, and occurrence probability is assessed against the baseline of z-values in a non-search space.

In the present study, the stimuli were padded by 100 ms of silence before and after the voice samples within noise to create a truly non-informative baseline search area of the acoustic signal ^68^. We set our thresholding to t > 2.5 and a probability of p = 0.05, following the threshold values (t > 2.3) recommended by Chauvin and colleagues ^71^. The resulting binary statistical classification images (0 non-significant, 1 significant) enable us to identify significant clusters that likely represent the spectrotemporal patches important for accurate affect classification per stimulus.

### Functional neuroimaging experiment (experiment 3)

We recruited an independent sample of 24 participants (15 female, mean age 26.4y, *SD* 3.6) via the same recruitment channels as described above. Structural and functional brain images were acquired on a Philips Ingenia 3T scanner using a standard 32-channel head coil. Due to the large number of trials, participants attended two separate scanning sessions. High-resolution structural scans were acquired at both sessions using a T1-weighted sequence (301 contiguous 1.2mm slices, repetition time (TR)/echo time (TE) = 1.96 s/3.71 ms, field of view 250mm, in-plane resolution 1×1mm). Functional images were partial volume acquisition using a multiband sequence (SENSE factor 3) based on a T2*-weighted echo-planar imaging (EPI) sequence covering the temporal and frontal cortex as well as subcortical structures (TR/TE 3290/30 ms, TA 980 ms, flip angle 88°, in-plane resolution 80×78mm; voxel size 1.71×1.71×3.9 mm; slice gap 0.3mm, 20 slices). A sparse sampling protocol was used so that each trial was presented without acquisition noise during 2310 ms silent gap between scans. Stimuli were presented between 1500-1790 ms after the onset of prior scan acquisition to avoid backward masking. Respiration and pulse rate were also acquired from each participant to correct for physiological artifacts during offline preprocessing.

In this experiment, stimuli were presented via Optoacoustics® OptoActive II active noise canceling headphones, canceling ∼20 dB of scanner noise and allowing comfortable and low background scanner noise levels without obstructing the ear canal. Stimuli were presented in eight blocks across the two scanning sessions, and each block had 15 catch trials including noise masks without any stimuli.

### Functional brain data preprocessing

Image preprocessing was conducted using SPM12 (version SPM12, www.fil.ion.ucl.ac.uk/spm/software/spm12). As the scanning took place over two sessions, functional images were realigned (motion corrected) and coregistered to the anatomical T1 acquisitions for their corresponding sessions. The second session T1 was then coregistered to the first session T1, and rigid-body transformation parameters were applied to all second session functional images. Images were spatially normalized to the Montreal Neurological Institute (MNI) stereotactic template brain using the T1 segmentation approach of Computation Anatomy Toolbox (CAT12; www.neuro.uni-jena.de/cat/), which provided normalization parameters. Functional images were transformed to MNI space using these normalization parameters and were then spatially smoothed using an isotropic Gaussian kernel of full-width-half-maximum of 9mm.

### Functional brain data modeling using a GLM approach

For the first-level analysis, we used the general linear model (GLM) to model stick functions for the onset of each individual trial, and we included 6 conditions in this GLM setup in a 3×2 design (3 emotions, 2 correct/uncorrected classification). Regressors of this emotion-by-correct setup were convolved with the HRF for each trial. Missed and catch trials were excluded from the model. Additional regressors of no interest were added for the six degrees of movement, heartbeat, and breathing to account for physiological noise in the data (TAPAS toolbox; https://www.tnu.ethz.ch/en/software/tapas). The six contrast images for each of the 6 conditions were then taken to a second-level group analysis.

For the second-level analysis, we used a fixed factorial design across the 6 conditions (3 emotions by correct/incorrect) and T-contrasts performed across dimensions of affect class or correct response. The group-level results were thresholded at a combined voxel threshold of *p<*0.005 and a cluster level of *k*=31. This combined voxel and cluster threshold corresponds to *p=*0.05 corrected at the cluster level and was determined by the 3dClustSim algorithm implemented in the AFNI software (https://afni.nimh.nih.gov/afni; version AFNI_18.3.01; including the new spatial autocorrelation function [ACF] extension) according to the estimated smoothness of the data across all contrasts.

### Reverse correlation analysis

The first-level analysis for the reverse correlation pipeline involved multiple processing stages (Fig. 4). We used the general linear model (GLM) to model stick functions for the onset of each 1200 individual trial separately on the unsmoothed images, now agnostic of trial condition or participant response. The stick function was again convolved with the HRF, resulting in beta-weight images on a trial-by-trial basis after model estimation. Additional regressors of no interest were again added for the six degrees of movement, heartbeat, and breathing to account for physiological noise in the data. The GLM was assessed for each participant individually, resulting in trial-by-trial beta-weight images for every participant. These images were advanced to the second level of the reverse correlation analysis to create correlation maps. Catch trials were not modeled in this analysis.

Next, the beta-weights per each brain voxel were split at the median brain activity level for this voxel into high-vs-low responding for all 200 trials per each of the 6 conditions. This resulted in 100 trials eliciting low activity (lower than median) and 100 trials eliciting high activity (higher than median) in this voxel for each condition. The index of high/low responding trials are used to build positive (mean of bubble masks from high responding trials, mean top 50%) and negative stimuli response maps (mean of bubble masks from low responding trials, mean bottom 50%) respectively. The negative response map was then subtracted from the positive for each of the 6 conditions. This process produced six difference maps (i.e., one for each condition) per each voxel and per each participant. Each of these difference response maps was then correlated (using a point biserial correlation analysis) to its corresponding general classification image per stimulus from the behavioral experiment (experiment 1). Correlation maps were smoothed with an isotropic Gaussian kernel of full-width-half-maximum of 9mm. Each participant was thus represented by six voxel-wise correlation maps that were taken into second-level group analysis.

We consider these behavior-brain correlation maps to be indicative of neural activity associated with the spectrotemporal patches revealed in the behavioral classification images for each stimulus. This analysis is so far agnostic of the participant’s performance in the task, and each voxel’s spectrotemporal response is only defined by the initial beta-weights and the indexing of high/low neural responding. Second-level group analysis was also conducted in SPM12 contrasts across the three affect classes, and statistical maps were thresholded at a voxel threshold of p=0.005 and a cluster size threshold of k=31, representing p=0.05 corrected at the cluster level.

### Linear mixed models for modelling ROI data

For the second-level group analysis, we performed a conjunction analysis (global null) and F-contrast to determine neural regions that show any common and differential effects between the affective voice classes, respectively. Significant regions from the conjunction and the F-contrast analysis were used to inform regions-of-interest (ROI) from which we extracted beta values across all 1200 trials for use in a linear mixed modeling (LME) approach. This allowed us to understand how correctness (correct trials, incorrect trials) was associated with single-trial brain signals in association with the ROI and affective voice class. The following equation was used for the LME modeling:

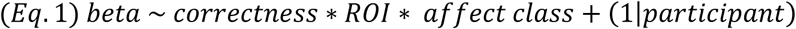

The ROIs were localized using the peak location (Fig. 6a-b). Using a 6mm spherical mask around the peak, we extracted mean voxel beta-weights for each single-trial beta image.

## AKNOWLEDGMENTS

The study was supported by the Swiss National Science Foundation (SNSF PP00P1_157409/1 and PP00P1_183711/1 to SF). We thank Jacqueline Te Heesen for her contribution to fMRI and behavioral data collection. We also thank Michael Mandel for his assistance and advice in creating bubbled auditory stimuli.

## AUTHOR CONTRIBUTIONS

HS contributed to designing the experiment, acquiring and analyzing the data, and writing the manuscript. LMA contributed to analyzing the data and writing the manuscript. SF contributed to designing the experiment, acquiring and analyzing the data, and writing the manuscript.

## DATA AVAILABILITY

The data that support the findings of this study are available from the corresponding authors upon reasonable request. No custom code was used in this study.

